# Conserved Roles for Receptor Tyrosine Kinase Extracellular Regions in Regulating Receptor and Pathway Activity

**DOI:** 10.1101/2020.05.29.122135

**Authors:** Monica Gonzalez-Magaldi, Jacqueline M. McCabe, Haley N. Cartwright, Ningze Sun, Daniel J. Leahy

## Abstract

Receptor Tyrosine Kinases (RTKs) comprise a diverse group of cell-surface receptors that mediate key signaling events during animal development and are frequently activated in cancer. Ligand-induced dimerization is the canonical mechanism by which RTKs are thought to be activated. We show here that deletion of the extracellular regions of 10 RTKs representing 7 RTK classes or their substitution with the dimeric immunoglobulin Fc region results in constitutive receptor phosphorylation but fails to result in phosphorylation of downstream signaling effectors Erk or Akt. Conversely, substitution of RTK extracellular regions with the extracellular region of the Epidermal Growth Factor Receptor (EGFR) results in increases in Erk and/or Akt phosphorylation in response to EGF. These results indicate that the activation signal generated by the EGFR extracellular region is capable of activating at least 7 different RTK classes. Failure of phosphorylated Fc-RTK chimeras to stimulate phosphorylation of downstream effectors indicates that either dimerization and receptor phosphorylation *per se* are insufficient to activate signaling or constitutive dimerization leads to pathway inhibition.

## Introduction

Receptor Tyrosine Kinases (RTKs) are Type I cell-surface proteins that consist of an extracellular ligand-binding region, a membrane-spanning helix, and a tyrosine kinase-containing intracellular region (Lemmon & Schlessinger, 2010). The human genome encodes 58 RTKs that assort into 20 classes based on homologous extracellular regions and cognate ligands. Distinct RTKs classes include receptors for Epidermal Growth Factor (EGF), Insulin (Ins), Fibroblast Growth Factors (FGFs), Nerve Growth Factor (NGF), Platelet-derived Growth Factor (PDGF), and Vascular Endothelial Growth Factor (VEGF). For typical RTKs, ligand binding to the extracellular region stimulates activity of the intracellular kinase and transphosphorylation of the receptor. Receptor phosphorylation results in recruitment of downstream effectors and initiation of signaling cascades that trigger changes in cell growth, differentiation, or behavior. RTK function is essential for normal development and maintenance of multicellular organisms, and abnormal RTK activity has been associated with birth defects and many cancers (Du & Lovly, 2018; McDonell, Kernohan, Boycott, & Sawyer, 2015). RTK-targeted therapies have proven effective treatments for many RTK-associated cancers, including colon, breast, and stomach cancers (Hynes & Lane, 2005; Roskoski, 2014; Yamaoka, Kusumoto, Ando, Ohba, & Ohmori, 2018).

The canonical mechanism by which RTKs are thought to act is ligand-induced receptor dimerization (Heldin, 1995; Heldin, Lu, Evans, & Gutkind, 2016; Yarden & Schlessinger, 1987). Several observations suggest that activation of RTK signaling is more complex than simple conversion of monomers to dimers, however. Firstly, specific deletions or mutations of RTK ECRs results in constitutive receptor phosphorylation indicating an autoinhibitory role for RTK ECRs in the absence of ligand (Arevalo et al., 2001; Ekstrand et al., 1994; Merlin et al., 2009; F. H. Qiu et al., 1988; Uren, Yu, Karcaaltincaba, Pierce, & Heidaran, 1997). Secondly, although ligand binding promotes dimerization of most RTKs and RTK extracellular regions *in vitro*, many RTKs dimerize in the absence of ligand (Clayton, Orchard, Nice, Posner, & Burgess, 2008; Del Piccolo & Hristova, 2017; Kozer et al., 2011; Liu et al., 2007; Macdonald & Pike, 2008; Martin-Fernandez, Clarke, Tobin, Jones, & Jones, 2002; Moriki, Maruyama, & Maruyama, 2001; Nagy, Claus, Jovin, & Arndt-Jovin, 2010; Saffarian, Li, Elson, & Pike, 2007; Zhang, Gureasko, Shen, Cole, & Kuriyan, 2006). Most notable in this respect are members of the Insulin Receptor (InsR) family, the subunits of which form disulfide-linked dimers and have long been thought to signal via a ligand-dependent conformational change (Lemmon & Schlessinger, 2010; Sparrow et al., 1997). Thirdly, in the case of the EGF Receptor (EGFR) artificially induced dimers result in receptor phosphorylation but not activation of downstream effectors (Liang et al., 2018; Yoshida et al., 2008), which suggests dimerization *per* se is insufficient to trigger pathway activation. Higher-order EGFR oligomers form in the presence of ligand and may be important for translating EGFR phosphorylation into pathway activation (Clayton et al., 2008; Gadella & Jovin, 1995; Huang et al., 2016; Kozer et al., 2013; Zanetti-Domingues et al., 2018).

To investigate the role of RTK extracellular regions (ECRs) in regulating RTK activity and the extent to which activation mechanisms are shared between different RTK classes, we examined the behavior of representatives of the EGFR, InsR, FGF Receptor (FGFR), VEGF Receptor (VEGFR), NGF Receptor (Trk), PDGF Receptor (PDGFR), and Met (Met and Ron) classes of RTK with deleted or substituted ECRs (Table 1). Early work had shown that EGFR/InsR chimeras retain function (Riedel, Dull, Honegger, Schlessinger, & Ullrich, 1989; Riedel, Dull, Schlessinger, & Ullrich, 1986), but such studies have not to our knowledge been extended beyond these two RTKs. We find that deletion or substitution of RTK ECRs with the dimeric immunoglobulin G Fc region generally results in constitutive phosphorylation of the receptor but failure to increase phosphorylation of the downstream effectors Erk or Akt. Conversely, substitution of ECRs with the EGFR ECR did not always result in detectable increases in receptor phosphorylation in response to EGF but did result in EGF-dependent increases in phosphorylation of Erk and/or Akt for all RTKs assayed. These results demonstrate that RTK ECRs generate a conserved activation signal and that constitutive phosphorylation of RTKs either leads to feedback inhibition of downstream effectors or an ECR generated signal in addition to dimerization is needed to couple receptor phosphorylation to pathway activation.

**Table 1.** Receptor Tyrosine Kinases Studied

| Name | Abbreviation | Ligand | Class |
| --- | --- | --- | --- |
| Epidermal growth factor receptor | EGFR | Human Epidermal growth factor | I |
| Insulin like growth factor receptor | IGF-1R | Human Insulin like growth factor | II |
| Insulin receptor | InsR | Recombinant Human Insulin Protein | II |
| Platelet-derived growth factor receptor alpha | PDGFRa | Human Platelet-Derived Growth Factor AA | III |
| Vascular endothelial growth factor receptor type 2 | VEGFR2 | Human Vascular Endothelial Growth Factor-121 | IV |
| Fibroblast growth factor type 1 and 2 | FDFR1/2 | Human Basic Fibroblast Growth Factor | V |
| Tropomyosin receptor kinase A | TrkA | Human β-Nerve Growth Factor | VII |
| Hepatocyte growth factor receptor | Met | Human Hepatocyte Growth Factor | X |
| Macrophage-stimulating protein receptor | Ron | Recombinant Human Macrophage Stimulating Protein | X |

## Results

### RTK ECRs prevent constitutive receptor phosphorylation

Deletions or mutations in several RTK extracellular regions have been associated with increased receptor phosphorylation, which has been interpreted as indicating an autoinhibitory role for these ECRs (Arevalo et al., 2001; Ekstrand et al., 1994; Merlin et al., 2009; F. H. Qiu et al., 1988; Shoelson, White, & Kahn, 1988; Uren et al., 1997). In most of these cases only a portion of the ECR is deleted, however. For example, the EGFR variant EGFRvIII, which is expressed in various cancer types, lacks roughly the N-terminal half of the ECR (aa 6-273) (Gan, Cvrljevic, & Johns, 2013). EGFRvIII is constitutively phosphorylated and able to activate downstream signaling pathways (Grandal et al., 2007; Huang et al., 2016; Schmidt, Furnari, Cavenee, & Bögler, 2003). Systematic deletion of RTK ECRs has not been carried out, however, and in isolated cases ECR deletion can lead to several-fold higher levels of receptor expression so that increased receptor phosphorylation may arise from misfolding during biogenesis or high receptor concentrations rather than loss of autoinhibition (Kavran et al., 2014).

To provide a systematic view of the role of RTK ECRs in receptor activation and downstream signaling, C-terminally HA-tagged variants of EGFR, InsR, IGF-1R, PDGFR, VEGFR, FGFR1, FGFR2, TrkA, Met and Ron with the entire ECR deleted (Figure 1) were transiently expressed in CHO cells and lysates from these cells analyzed 24 hours after transfection by Western blot. Although phosphorylation of PDGFRα was relatively low, all ΔECR variants proved constitutively phosphorylated, consistent with an autoinhibitory function for the ECRs (Figure 2A and 2B).

**Figure 1.**
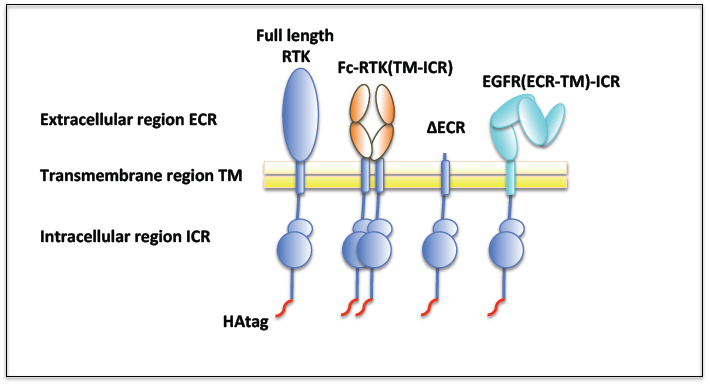
Schematic diagram of chimeric receptors. Cartoon representations and abbreviations for each of the native or variant RTKs used in this study are shown. In addition to full-length native RTKs, which are composed of extracellular, transmembrane, and intracellular regions, variants in which the extracellular region was replaced with a constitutively dimeric murine IgGFc (Fc-RTK(TM-ICR)), the extracellular region was deleted (ΔECR), or the extracellular and transmembrane regions replaced with the human EGFR extracellular and transmembrane regions (EGFR(ECR-TM)-ICR) were created. Abbreviations for specific RTKs shown in Table 1 are substituted in the above designations to denote specific RTK variants in the text. A C-terminal HA tag added to each variant to aid uniform detection and is shown in red.

**Figure 2.**
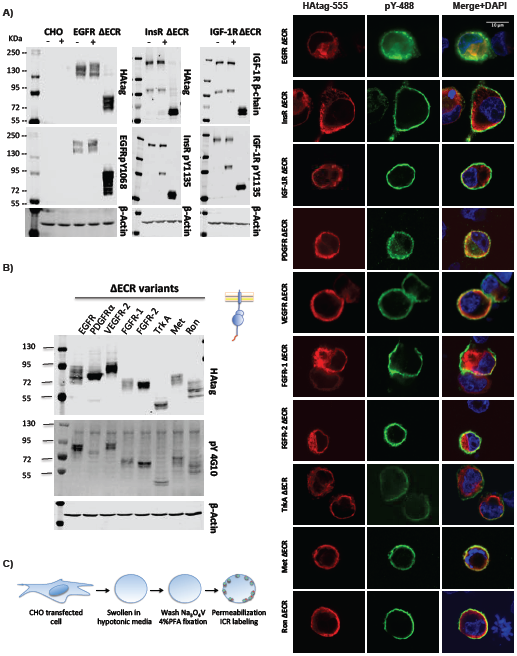
ΔECR variant RTKs are constitutively phosphorylated and trafficked to the plasma membrane. Western blot analysis of expression (HA-tag) and phosphorylation (EGFR pY1068, InsR/IGF-1R pY11135, or pY 4G10) of transiently-transfected CHO cells expressing HA-tagged versions of **A)** full length and ΔECR forms of EGFR, InsR, and IGF-1R with and without addition of cognate ligands, **B)** ΔECR-RTK variants of EGFR, PDGFRα, VEGFR-2, FGFR-1, FGFR-2, TrkA, Met, and Ron. β-Actin was included as a loading control, and the same amount of total protein was loaded in each well. Representative Western blots from three independent experiments are shown. **C)** Confocal microscopy images of CHO cells transfected with the indicated ΔECR RTK variant showing cell surface expression and phosphorylation for each variant. Cells were swollen in hypotonic media, washed, permeabilized and stained with an anti-HA antibody as indicated in the cartoon (HAtag-555; red), an anti-phosphotyrosine antibody (pY-488; green), and DAPI (nuclei; blue). The scale bar represents 10 μm. Non-transfected cells evident with DAPI in the same field serve as negative controls for primary and secondary antibodies. See also Figure S1 and S2.

To determine whether ΔECR variants are trafficked to the cell surface and rule out that the observed phosphorylation arises from aggregation in intracellular compartments, cells expressing each ΔECR variant were swollen, permeabilized, and stained with antibodies against the HA-tag and phosphotyrosine. In each case, including PDGFRα, phosphorylated forms of ΔECR variants were observed at the cell surface indicating that misfolding or aggregation during biogenesis cannot wholly account for phosphorylation of ΔECR variants (Figure 2C). Colocalization of the HA-tagged receptor ΔECR variants with cellular lectins in permeabilized cells as judged by immunofluorescence (Figure S1A) and a cell-surface biotinylation assay (Figure S1B) for all the ΔECR variants with an extracellular FLAG tag confirmed their plasma membrane localization.

### Transiently expressed native RTKs traffic to the cell surface but have unprocessed intracellular forms

Full-length native forms of each studied RTK bearing C-terminal HA-tags were expressed in CHO cells and stimulated with 100 and 500 ng/ml of their cognate ligand (Figure S2A). Cell lysates were then assayed by Western blot for receptor expression and phosphorylation. In addition to receptor forms migrating in polyacrylamide gels at the expected molecular weights for glycosylated receptors, faster migrating and constitutively phosphorylated receptor forms with molecular weights consistent with unglycosylated or incompletely glycosylated receptors were observed for all receptors except Ron. Treatment of cell cultures with Tunicamycin and cell lysates with PNGaseF collapsed slower migrating bands into these faster migrating bands (Figure S2B), and surface biotinylation experiments demonstrated enrichment of slower-migrating, glycosylated receptor forms at the cell surface (Figure S2C). The faster migrating bands thus reflect intracellular accumulation of unglycosylated or incompletely glycosylated receptor forms. Based on this observation, ligand-dependent phosphorylation was only measured for slower migrating receptor forms. Contrary to expectation, addition of increasing concentrations of cognate ligands resulted in only modest increases in phosphorylation of the glycosylated receptor forms (Figure S2A), but, as shown below, resulted in increased phosphorylation of downstream effectors indicating functional receptors and ligands.

### Fc-driven but not EGFR-ECR-driven RTK dimerization leads to robust receptor phosphorylation

To assess whether dimerization *per se* is sufficient for RTK activation or whether specific types of dimers are needed, chimeric receptors were created in which the ECRs of InsR, IGF-1R, PDGFRα, VEGFR, TrkA, FGFR1, FGFR2, Met, and Ron were substituted with (i) a constitutively dimeric immunoglobulin G constant region (Fc) or (ii) the EGFR extracellular and transmembrane regions (ECR-TM) (Figure 1). We have previously shown using FRET microscopy that substituting the EGFR ECR with Fc results in constitutive cell-surface dimers (Byrne, Hristova, & Leahy, 2020).

Fc-RTK chimeras with the Fc region fused to the EGFR, IGF-1R, PDGFRα, VEGFR, FGFR1, FGFR2, TrkA, Met, and Ron transmembrane and intracellular regions expressed well, reached the cell surface as judged by immunofluorescence (Figure S3), and were constitutively phosphorylated as judged by immunofluorescence and Western blot (Figure 3A). The Fc-InsR chimera expressed at lower levels and was less phosphorylated relative to other Fc chimeras. Even though each of the chimeric receptors with the EGFR ECR (ECR-TM) was expressed and reached the cell surface as judged by immunofluorescence (Figure S4), a clear increase in chimeric receptor phosphorylation in response to addition of EGF was only observed by Western blot for the InsR, IGF-1R, FGFR1, FGFR2, and TrkA chimeras (Figure 3B). Owing to ambiguity concerning statistical analysis of triplicate results we have opted to present band intensities of treated conditions normalized to the matched untreated condition (Figure 3B). The data points from each of three independent experiments were then plotted to enable direct visualization of the amplitudes and spread of observed changes.

**Figure 3.**
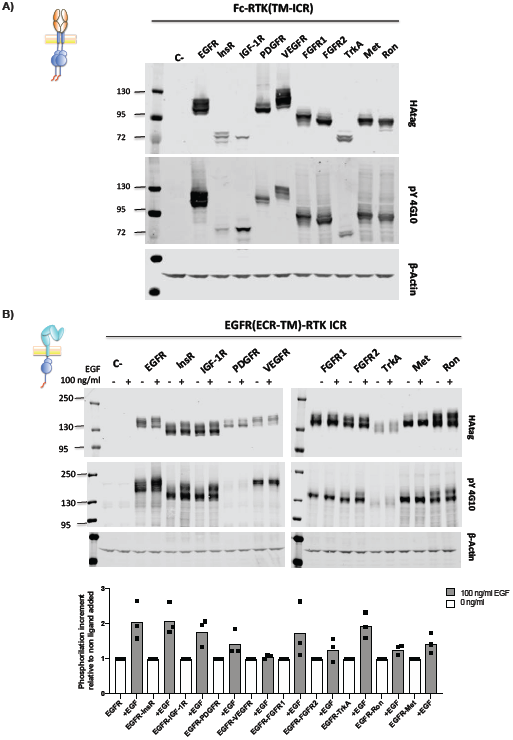
All Fc-driven but not all EGFR-ECR-driven RTK dimerization leads to receptor phosphorylation. Chimeric receptors with **A)** the IgGFc domain (Fc-RTK(TM-ICR) or **B)** the EGFR extracellular and transmembrane regions (EGFR(ECR-TM)-ICR) substituted for the native ECR or ECR and transmembrane region, respectively, were transiently expressed in CHO cells and cell lysates analyzed by Western blot for expression (HA-tag) and phosphorylation (pY4G10). In the case of EGFR(ECR-TM)-ICRs, 100 ng/ml of EGF was added for 5 min before lysing the cells. All Fc-RTK receptor variants were constitutively phosphorylated, but, with the exception of EGFR-InsR, EGFR-IGF-1R, FGFR1, FGFR2 and TrkA chimeras, phosphorylation of EGFR(ECR-TM)-ICR variants chimeras did not increase notably when ligand was added. The graphic representation of the increment in phosphorylation for each chimeric receptor relative to the non-ligand added condition is shown. Dots represent an independent experiment for each condition and bars represent the mean value of the three experiments. See also Figures S3 and S4.

### EGFR-ECR-activated but not Fc- or Δ ECR-activated RTKs result in robust phosphorylation of downstream effectors

RTK phosphorylation is frequently used as a proxy for RTK signaling (Lemmon & Schlessinger, 2010; Pawson, 2004), but it has been shown that phosphorylation of artificially dimerized EGFR does not necessarily lead to phosphorylation of downstream effectors (Liang et al., 2018; Sousa et al., 2012). To determine if phosphorylation of EGFR(ECR-TM)-RTK(ICR), Fc-RTK(TM-ICR), and ΔECR-RTK(TM-ICR) variants leads to phosphorylation of downstream effectors, Western blots for Erk, Akt, phospho-Erk, and phospho-Akt were carried out for each RTK variant and for native full-length receptors in the absence and presence of ligand (Figures 4 and 5). In all cases, addition of the cognate ligand to the native receptor led to increased phosphorylation of Erk, Akt, or both (Figure 4). VEGFR expression was much lower than other RTKs, however, which may explain the more modest increases in Erk and Akt phosphorylation seen following treatment of VEGFR-expressing cells with VEGF (Figure 4). Owing to ambiguity concerning the significance of statistical analysis of triplicate results from Western blot experiments performed on multiple gels and best way to present results, we have opted to normalize integrated band intensities of treated conditions to those of untreated conditions and present these values relative to the treated condition for native EGFR. The data points for each experiment are plotted to enable direct visualization of the relative amplitudes and spread of observed changes (Figure 4B and Figure 5B).

**Figure 4.**
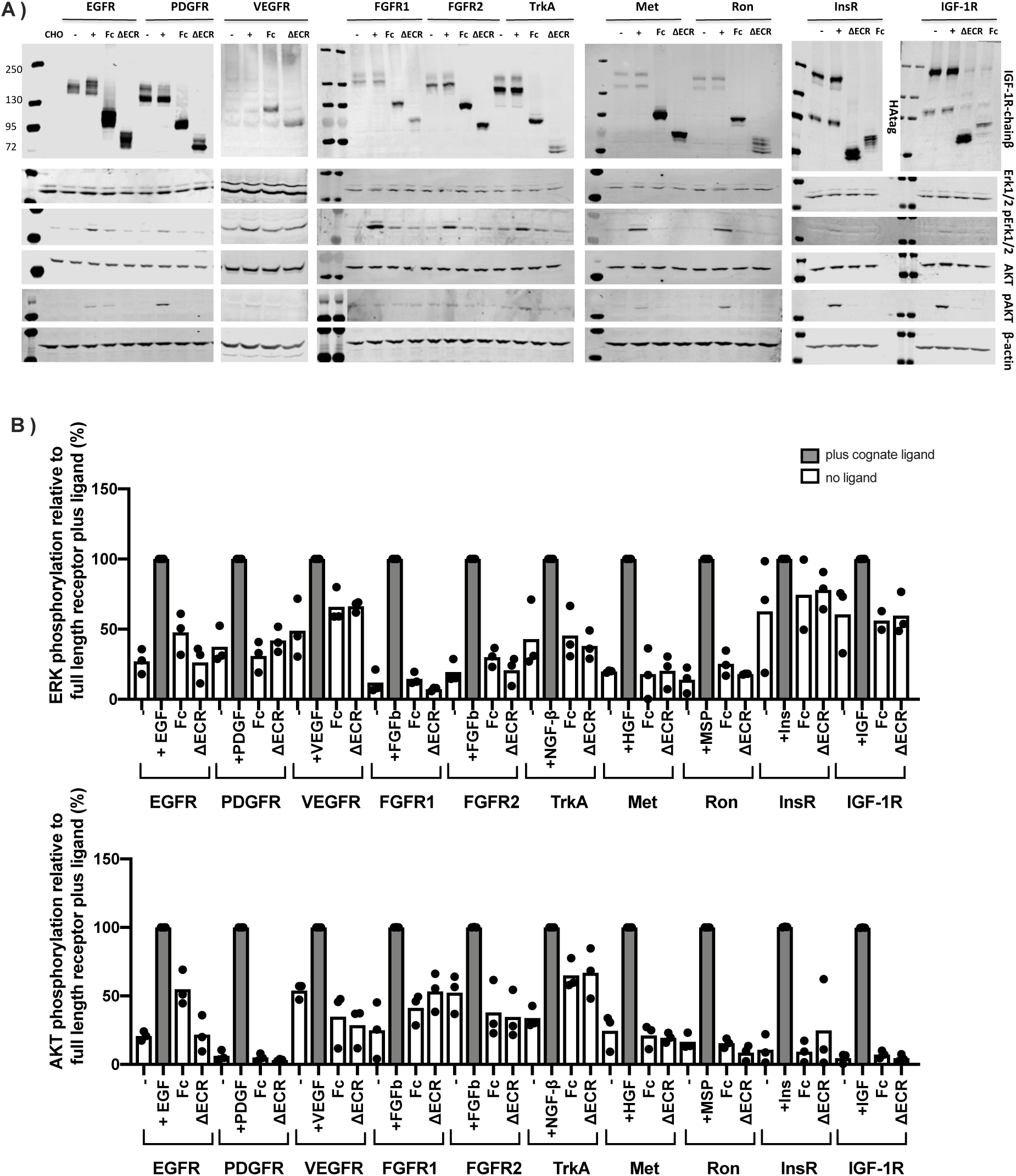
RTKs(ECR)-activated but not Fc- or ΔECR-phosphorylated RTKs trigger increased phosphorylation of downstream effectors. **A)** Western blot analysis of RTK variant expression (HA-tag or IGF-1R β-chain), Erk 1/2 expression (Erk 1/2), phospho-Erk 1/2 (pErk1/2), Akt expression (Akt), and phospho-Akt (pAkt) levels for native RTKs with and without cognate ligand, Fc-RTK(TM-ICR) (Fc), and ΔECR (ΔECR) variants of the indicated RTKs transiently expressed in CHO cells. β-Actin was included as a loading control, and the same amount of total protein was loaded in each well. **B)** Quantification and graphic representation of the downstream activation obtained from WB analysis. Each dot represents an independent experiment for each condition, and the bar heights represent the mean values for these experiments.

**Figure 5.**
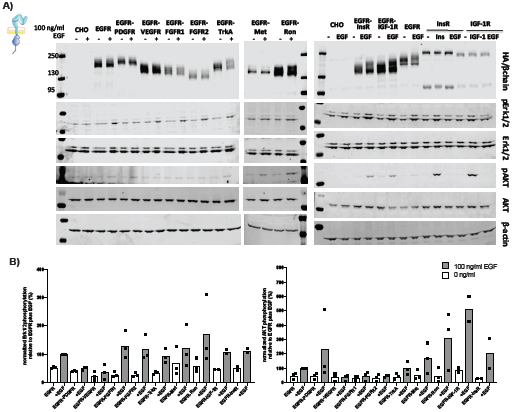
EGFR(ECR-TM)-RTK chimeras activate downstream pathways in response to EGF. **A)** Western blot analysis of cell lysates from CHO cells transiently expressing the indicated chimeric RTKs in which the EGFR ECR-TM region was substituted for the native ECR-TM with and without EGF (or cognate ligands) are shown. Chimeric receptor EGFR(ECR-TM)-RTK expression (HA-tag or IGF-1R β-chain), phospho-Erk 1/2 (pErk1/2), Erk 1/2 expression (Erk 1/2), phospho-Akt (pAkt), and Akt expression (Akt) were assessed. β-Actin was included as a loading control, and the same amount of total protein was loaded in each well. **B)** Quantification and graphic representation of phospho-Erk and phospho-Akt levels obtained from WB analysis normalized as detailed in results. Each dot represents an independent experiment, and the bar heights represent the mean values for three experiments (except for pERK for Fc-InsR and Fc-IGF-1R, which have two).

Despite constitutive phosphorylation of the ΔECR RTKs (Figure 2B), no ΔECR RTK variant stimulated detectable increases in phosphorylation of Erk or Akt (Figure 4A). Similarly, no Fc-RTK chimera except Fc-EGFR stimulated detectable phosphorylation of Erk or Akt (Figure 4). Fc-EGFR was the most highly expressed Fc chimera, however, and only modest levels of phospho-Erk and phospho-Akt relative to wild-type EGFR in the presence of EGF were observed (Figure 4). In contrast, clear EGF-dependent phosphorylation of Erk, Akt, or both was observed for all EGFR-ECR chimeras except the VEGFR chimera, for which Erk phosphorylation was present but modest and no increase in Akt phosphorylation was observed (Figure 5). This observation is somewhat surprising given the undetectable to modest phosphorylation of many chimeras themselves when stimulated with EGF (Figure 3B).

The absence of phosphorylation of downstream effectors in cells expressing constitutively phosphorylated Fc-chimera and ΔECR RTK variants may stem from inhibitory feedback mechanisms induced by constitutive receptor phosphorylation. To determine whether Erk and Akt signaling was generally repressed in cells expressing these RTK variants, CHO cells were co-transfected with wild type EGFR and Fc-EGFR or ΔECD-EGFR in ratios that resulted in similar expressions levels. In these cells both full length EGFR and downstream effectors became phosphorylated in response to EGF as judged by Western blot despite the presence of constitutively phosphorylated Fc-EGFR or ΔECR-EGFR variants (Figure S5). This observation indicates that Fc-EGFR and ΔECR-EGFR do not induce generalized inhibition of EGFR signaling.

## Discussion

The canonical mechanism by which ligands are thought to activate RTKs is by formation of ligand-dependent receptor dimers (Heldin, 1995; Heldin et al., 2016; Yarden & Schlessinger, 1987). Although ligands drive dimerization of most RTKs (Lemmon & Schlessinger, 2010), several observations imply that regulation of receptor activity is more complex than simple conversion of monomers to dimers (Clayton et al., 2008; Del Piccolo & Hristova, 2017; Huang et al., 2016; Zanetti-Domingues et al., 2018). Using the well-studied EGFR as an example, preformed and presumably inactive EGFR dimers are present in the absence of ligand (Byrne et al., 2020; Freed, Alvarado, & Lemmon, 2015; Gadella & Jovin, 1995; Macdonald & Pike, 2008; Macdonald-Obermann, Adak, Landgraf, Piwnica-Worms, & Pike, 2013; Sako, Minoghchi, & Yanagida, 2000; Tao & Maruyama, 2008), higher-order EGFR oligomers appear in the presence of ligand (Gadella & Jovin, 1995; Huang et al., 2016; Kozer et al., 2013; Zanetti-Domingues et al., 2018), and dimerization-driven phosphorylation of EGFR can be insufficient to activate downstream signaling effectors (Liang et al., 2018; Yoshida et al., 2008).

To investigate regulatory roles of RTK ECRs beyond mediating ligand-dependent dimers and whether these roles are shared among diverse RTKs, we examined the behavior of variant forms of 10 RTKs (EGFR, InsR, IGF-1R, PDGFRα, VEGFR, FGFR1, FGFR2, TrkA, Met, and Ron) representing 7 RTK classes. Deletion of the ECR of each RTK led to expression of a constitutively phosphorylated receptor at the cell surface consistent with a general autoinhibitory role for RTK ECRs. Increased expression levels of the ΔECR variants relative to native receptors cannot be ruled out as contributing to this phosphorylation, however. The failure of any ΔECR variant to stimulate phosphorylation of Erk or Akt despite their overexpression and relatively high phosphorylation levels implies that either ligand-bound RTK ECRs provide a signal in addition to dimerization that is needed to couple receptor phosphorylation to pathway activation or constitutive receptor phosphorylation induces feedback inhibition of downstream effectors. If an additional signal is present, the presence of Erk phosphorylation in cells expressing the oncogenic EGFRvIII (Pedersen et al., 2005; Pedersen, Tkach, Pedersen, Berezin, & Poulsen, 2004), which lacks the first two of four EGFR extracellular domains, may provide a clue to its nature. Mutations in a region in Domain IV of the EGFR ECR, which is present in EGFRvIII but not in ΔECR EGFR, have been shown to decrease EGFR activity as well as formation of higher-order EGFR oligomers (Huang et al., 2016) and this region of Domain IV may confer an activity needed to activate downstream effectors.

Although all Fc-RTK chimeras were highly expressed relative to native receptors, trafficked to the cell surface, and constitutively phosphorylated, none but Fc-EGFR led to detectable increases in phosphorylation of Erk or Akt in conditions where much lower levels of native receptors led to increased phosphorylation of Erk and/or Akt in the presence of ligand. Fc-EGFR was the most overexpressed Fc-RTK chimera but only resulted in weak phosphorylation of Erk and Akt relative to ligand-stimulated EGFR, consistent with an impaired ability of Fc-EGFR to trigger Erk or Akt activation. This result may indicate that while Fc-mediated dimerization is sufficient to result in constitutive receptor phosphorylation it is not sufficient to trigger phosphorylation of downstream effectors. Induction of pathway inhibition by constitutive phosphorylation cannot be ruled out as underlying the absence of Erk and Akt phosphorylation, but the ability of native EGFR to activate downstream effectors in response to ligand when co-transfected with Fc-EGFR indicates that any induced pathway inhibition is not global. This observation coupled with remarkable earlier observations that artificially induced EGFR dimerization also led to receptor phosphorylation but not activation of downstream effectors raises the possibility that ligand-bound EGFR ECRs provide a signal in addition to dimerization that is essential to trigger pathway activation (Liang et al., 2018; Yoshida et al., 2008).

Conversely, although addition of EGF to EGFR (ECR-TM)-RTK chimeras did not lead to detectable increases in phosphorylation for several chimeras, increased phosphorylation of Erk and/or Akt was observed following addition of EGF to each of the EGFR(ECR-TM)-RTK(ICR) chimeras. The ability of the EGFR ECR to supply this role for at least 6 additional classes of RTKs indicates that a common signaling mechanism is shared among most if not all signaling-competent RTK ECRs. Given that dimerization *per se* may not be sufficient to trigger RTK pathway activation, it is tempting to speculate what additional shared signal may be required to trigger pathway activation. As higher-order EGFR oligomers are known to form in the presence of ligand, an intriguing possibility is that such oligomers might constitute part of this additional signal. In addition to any autoinhibitory mechanism supplied by ECRs in the absence of ligand, the need for a higher-order RTK oligomer to trigger pathway activation would minimize pathway activation from random collisions in the cell membrane. Higher-order oligomers could also contribute to downstream signaling by increasing the local concentration and stability of adaptors recruited to phosphorylated RTKs as well as potentially altering local features of the cell membrane (Ichinose, Murata, Yanagida, & Sako, 2004; Pawson, 2004).

Collectively, the results reported here confirm and generalize an autoinhibitory role for RTK ECRs, demonstrate that the signal generated by the ligand-bound EGFR ECR is sufficient to activate members of at least seven RTK families, and hints that, as observed by others for EGFR (Liang et al., 2018; Yoshida et al., 2008), a signal in addition to dimerization may be needed by most if not all RTKs to couple receptor phosphorylation to activation of downstream effectors. Future work will be needed to confirm the presence of this additional signal and, if present, define its precise nature, whether other cellular factors are involved, and its role in RTK-associated cancers and RTK-targeted anticancer therapies.

## Acknowledgements

This work was supported by NIH R01GM099321 (DJL) and CPRIT RR160023 (DJL). We thank Pat Byrne and other Leahy lab members for advice and commentary, Pat Byrne for providing the FcEGFR clone, and Kalina Hristova for providing the VEGFR, FGFR and TrkA clones.

## Author Contributions

Conceptualization, M.G-M., J.M.M., and D.J.L.; Methodology, M.G-M., J.M.M., and D.J.L.; Investigation, M.G-M., J.M.M., H.N.C., and N.S.; Writing–Original Draft, D.J.L. and M.G-M.; Writing–Review & Editing, D.J.L. and M.G-M.; Funding Acquisition, D.J.L.

## Declaration of Interests

The authors declare no competing interests.

## Materials and Methods

Key Resources Table

| **Key Resources Table** |  |  |  |  |
| --- | --- | --- | --- | --- |
| **Reagent type (species) or resource** | **Designation** | **Source or reference** | **Identifiers** | **Additional information** |
| gene | Full length human IGF-1R |  | NM_000875.4 |  |
| gene | Full length human InsR |  | NM_000208.2 |  |
| gene | Full length human EGFR |  | NM_005228.3 |  |
| gene | Full length human PDGFRa |  | NM_006206 |  |
| gene | Full length human VEGFR2 |  | NM_002253.2 |  |
| gene | Full length human FGFR1 |  | NM_023110.2 |  |
| gene | Full length human FGFR2 |  | NM_000141.4 |  |
| gene | Full length human TrkA |  | NM_001012331.1 |  |
| gene | Full length human Ron |  | NM_002447.2 |  |
| gene | Full length human Met |  | NM_000245.2 |  |
| gene | IgG Fc mouse |  | U65534 |  |
| cell line | Chinese hamster ovary cells CHO-K1 | ATCC | Cat# CCL-61 (RRID: CVCL_0214) |  |
| antibody | Mouse anti-Phosphotyrosine clone 4G10 | EMD Millipore | Cat # 05-321 (RRID: AB_568857) | WB(1:100)<br>IF(1:200) |
| antibody | Rabbit anti-HA Polyclonal antibody | Thermo Fisher Scientific | Cat # PA1-985 (RRID: AB_1958085) | WB(1:100)<br>IF(1:200) |
| antibody | Rabbit anti phospho-EGFR pTyr1068 antibody | Thermo Fisher Scientific | Cat # 44-788G (RRID: AB_2533754) | Dilution 1:1000 |
| antibody | Rabbit anti-EGFR D1D4J | Cell Signaling Technology | Cat # 54359 (RRID: AB_2799458) | Dilution 1:200 |
| antibody | Rabbit anti IGF-1R β-chain D23H3 | Cell Signaling Technology | Cat # 9750 (RRID: AB_2797674) | Dilution 1:1000 |
| antibody | Rabbit anti Erk1/2 MAPK p42/p44 | Cell Signaling Technology | Cat # 4695 (RRID: AB_390779) | Dilution 1:1000 |
| antibody | Rabbit anti pErk P-p42/p44 MAPK | Cell Signaling Technology | Cat # 9101 (RRID: AB_331646) | Dilution 1:1000 |
|  | (Erk1/2)<br>Thr202/Tyr<br>204 |  |  |  |
| antibody | Rabbit anti-AKT1 | Sigma-Aldrich | Cat # SAB4300575<br>(RRID: AB_10624863) | Dilution<br>1:1000 |
| antibody | Rabbit anti-<br>β-Actin | Cell<br>Signaling<br>Technolo<br>gy | Cat # 4968 (RRID:<br>AB_2313904) | Dilution<br>1:2000 |
| antibody | Secondary<br>Goat anti-<br>rabbit-680RD | Li-Cor | Cat # 926-68071 | Dilution<br>1:15000 |
| antibody | Secondary<br>Goat anti-<br>mouse<br>IgG<sub>2b</sub>-<br>800CW | Li-Cor | Cat # 926-32352 | Dilution<br>1:15000 |
| antibody | Rabbit anti-Phospho-Akt<br>(Ser473) | Cell<br>Signaling<br>Technolo<br>gy | Cat # 4060 (RRID:<br>AB_2315049) | Dilution<br>1:1000 |
| antibody | Rabbit anti-Hsp90α | Cell<br>Signaling<br>Technolo<br>gy | Cat # 4877 (RRID:<br>AB_2233307) | Dilution<br>1:1000 |
| antibody | DYKDDDD<br>K Tag<br>Polyclonal<br>Antibody | Thermo<br>Fisher<br>Scientific | Cat # PA1-984B (RRID:<br>AB_347227) | Dilution<br>1:2000 |
| antibody | Goat anti<br>rabbit IgG<br>Alexa fluor<br>555 | Thermo<br>Fisher<br>Scientific | Cat # A-21428 (RRID:<br>AB_2535849) | Dilution<br>1:500 |
| antibody | Donkey anti<br>mouse IgG<br>Alexa fluor<br>555 | Invitrogen | Cat # A-31570 (RRID:<br>AB_2536180) | Dilution<br>1:500 |
| antibody | Rabbit anti<br>phospho-IGF-I<br>receptor $\beta$<br>Tyr1135/36/<br>IR $\beta$ | Cell<br>Signaling<br>Technolo<br>gy | Cat # 2969 (RRID:<br>AB_11178660) | Dilution<br>1:1000 |
|  | Tyr1150/1151 |  |  |  |
| recombinant DNA reagent | vector pαH | Aricescu et al. 2006 | N/A |  |
| recombinant DNA reagent | pcDNA 3.1 | Invitrogen | Cat # V79020 |  |
| peptide, recombinant protein | Human Epidermal growth factor EGF | (C. Qiu et al., 2009) | N/A |  |
| peptide, recombinant protein | Human Insulin like growth factor IGF-1 | (Kavran et al., 2014) | N/A |  |
| peptide, recombinant protein | Recombinant Human Insulin Protein | Novus Biologicals | Cat # NBP1-99193 |  |
| peptide, recombinant protein | Human Platelet-Derived Growth Factor AA | Cell Signaling | Cat # 8913 |  |
| peptide, recombinant protein | Human Basic Fibroblast Growth Factor | Cell Signaling | Cat # 8910 |  |
| peptide, recombinant protein | Human Vascular Endothelial Growth Factor-121 | Cell Signaling | Cat # 8908 |  |
| peptide, recombinant protein | Human β-Nerve Growth Factor | Cell Signaling | Cat # 5221 |  |
| peptide, recombinant protein | Human Hepatocyte Growth Factor | Sigma-Aldrich | Cat # H9661 |  |
| peptide, recombinant protein | Recombinant Human Macrophage Stimulating Protein | R&D systems | Cat # 352-MS-010 |  |
| commercial assay or kit | Pierce™ Cell Surface Biotinylation and Isolation Kit | Pierce | Cat # A44390 |  |
| commercial assay or kit | Pierce™ BCA Protein Assay | Thermo Fisher Scientific | Cat # 23225 |  |
| chemical compound, drug | Wheat Germ Agglutinin Alexa Fluor 488 conjugate | Invitrogen | Cat # W11261 |  |
| chemical compound, drug | DAPI (4',6-Diamidino-2-Phenylindole, Dihydrochloride) | Thermo Fisher Scientific | Cat # 62247 |  |
| chemical compound, drug | ProLong™ Diamond antifade mountant | Thermo Fisher Scientific | Cat # P36965 |  |
| software, algorithm | Fiji | [Fiji contributors](https://imagej.net/Fiji/contributors) | https://imagej.net/Fiji/Downloads |  |
| software, algorithm | Prism Graphpad | GraphPad | https://www.graphpad.com/scientific-software/prism/ |  |
| software,<br>algorithm | Zen lite<br>software | Zeiss | https://www.zeiss.com/microscopy/us/products/microscope-software/zen-lite.html |  |
| software,<br>algorithm | Image<br>studio™ lite<br>software | Li-Cor | https://www.licor.com/bio/image-studio-lite/download |  |

### Cell culture, Transfection and Expression

CHO-K1 cells were maintained in adherent culture in DMEM: F12 (Gibco) supplemented with 5% Fetal Bovine Serum (Gibco). For transient transfections, CHO-K1 cells were plated in six-well plates at 0.5 × 10^6^ cells/well and transfected with 1 μg of each indicated expression plasmid using Lipofectamine 2000 (Thermo Fisher Scientific) according to the manufacturer’s protocol. After 18 hours cells were washed three times with 2 ml Ham’s F12 supplemented with 1 mg/ml BSA and serum starved in this medium for 3 hr at 37°C. Each specific ligand was added in designated wells, 100 ng/ml EGF for 5 min, 150 ng/ml IGF1, 200 nM (1.14 μg/ml) insulin for 30 min, 100 ng/ml PDGF-AA, 100 ng/ml FGF-basic, 100 ng/ml hVEGF-121, 100 ng/ml Human β-Nerve Growth Factor, 100 ng/ml Hepatocyte Growth Factor, 100 ng/ml Macrophage Stimulating Protein and incubated 15 min at 37°C. Wells were washed with ice-cold phosphate buffered saline and lysed for 30 min at 4°C in 250 μl of RIPA buffer (50 mM Tris pH 8.0, 150 mM NaCl, 1% (v/v) NP40, 0.5% (w/v) sodium deoxycholate, 0.1% (w/v) sodium dodecyl sulfate) supplemented with 1 mM activated Na_3_VO_4_, Pierce protease inhibitor minitablet (Thermo Fisher Scientific), Benzonase nuclease (Sigma), and 10 nM iodoacetamide to prevent disulphide-bond formation during lysis. The total protein concentration of clarified lysates was determined using the BCA assay (Smith et al., 1985), and all lysates were adjusted to lowest total protein concentration using RIPA buffer.

Cotransfection experiments were performed in the same growth conditions, with the ratio of DNAs encoding different variants adjusted to obtain similar levels of expression for both variants. For the pair EGFR:ΔECR-EGFR the DNA ratio was 1:1 with 1.5 μg of each expression plasmid transfected using 1mg/ml PEI as transfecting agent as described (Longo, Kavran, Kim, & Leahy, 2013) (*3:1 ratio of PEI to DNA (w/w)*). For the EGFR:Fc-EGFR pair the ratio was 2:0.5. Cells were lysed 20 h post transfection following serum starvation described above.

### Expression vector design

DNA sequences encoding each RTK variant were cloned into modified versions of the expression vector pαH, which was derived from pHLSec by insertion of an alternative multiple cloning site (Aricescu, 2006). Modifications to pαH included: removal of the signal sequence encoding region, addition of a region encoding a C-terminal HA tag, and addition of coding sequences for the EGFR (ECR-TM) that facilitated cloning of chimeric receptors.

DNA sequences encoding full length human IGF-1R, InsR, EGFR VEGFR2, PDGFRα, FGFR1 and FGFR2, TrkA, Ron, and Met cDNAs were used to generate full length and chimeric receptors. Deletions of the ECR for each receptor were generated by PCR. Fc-RTK chimeric receptors substituted regions encoding the mouse IgG Fc region for native ECRs prior to the TM-ICR RTK of interest (Table 2). Transmembrane boundaries for each RTK were guided by domain annotation of mRNA and protein sequences from NCBI.

**Table 2.**
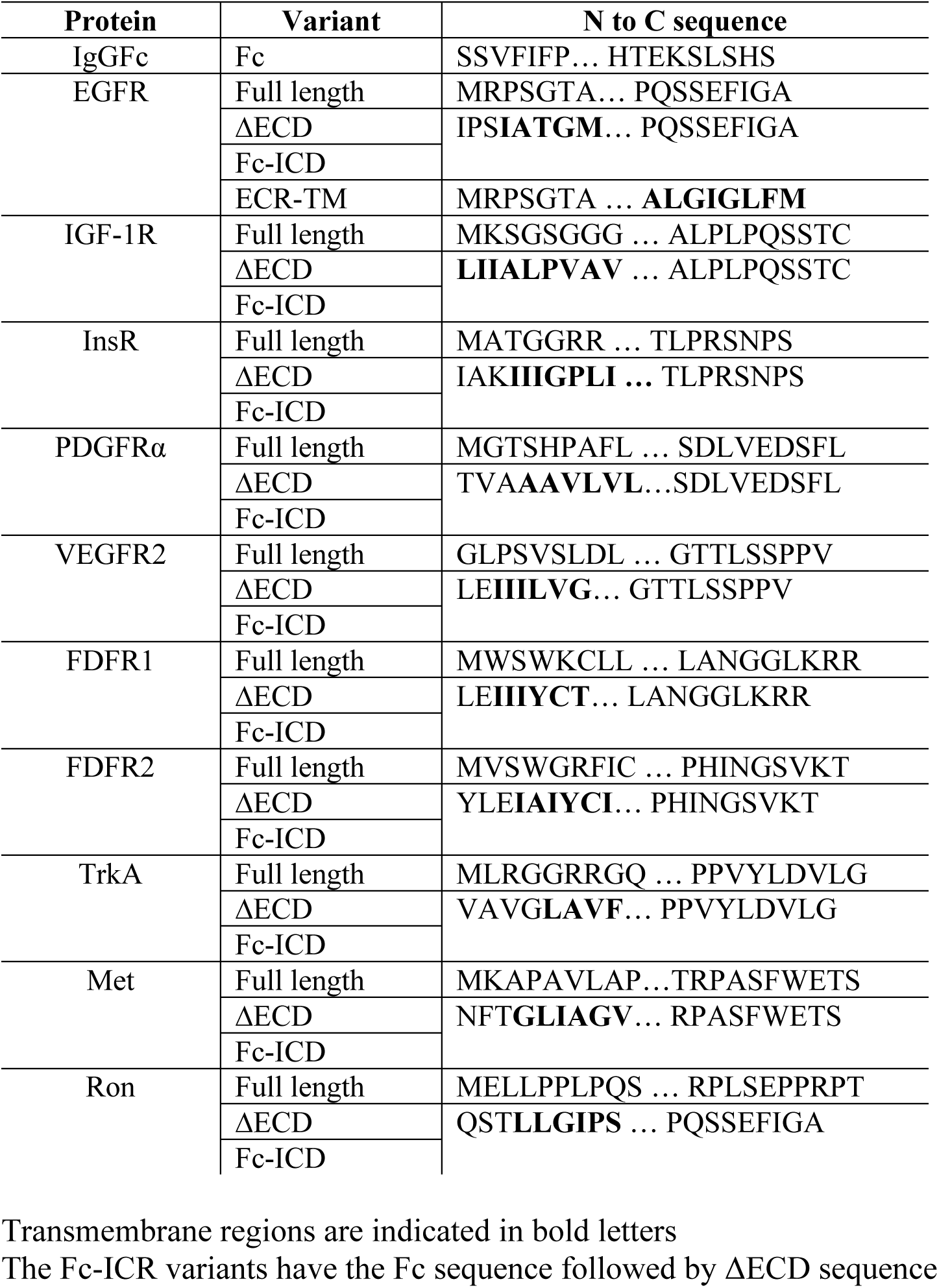
Amino acid sequences of RTK variant termini and junctions

### Western Blots

Cell lysates with normalized total protein concentrations were mixed with SDS sample buffer and boiled, separated by SDS-PAGE (Novex 4-12% Tris-Glycine Mini Protein Gels, 1.0mm, 15 well (Life Technologies, EC6025)), and transferred using the iBlot® Dry Blotting System with iBlot® Transfer Stacks to a nitrocellulose membrane (Life Technologies, IB301001). The membrane was blocked with 3% low fat milk in TBS, and proteins were labeled by incubation with the selected primary antibody and the corresponding Infrared dye secondary antibody, which was detected using a Li-Cor Odyssey Clx Near IR imaging system. The amount loaded in each well was normalized by BCA assay. β-actin was used as loading control as a qualitative confirmation of overall protein abundance present in each experiment. Band intensities were integrated using Image Studio Lyte software. Independent experiments for each condition are presented as dots in the graphic representations with bar heights representing the mean value of multiple measurements. Statistical analyses are not reported owing to ambiguity interpreting the meaning of statistics for triplicate experiments. We present instead the plots showing data points from each experiment and their mean.

### Immunofluorescence and confocal microscopy

CHO-K1 cells grown on glass cover slips were transfected with 1 μg of the indicated DNA. At 24 hours post transfection, cells were fixed in 4% paraformaldehyde for 15 min at room temperature, blocked and permeabilized with PBTG buffer (0.1% Triton X-100, 1% bovine serum albumin (BSA), and 1 M glycine in PBS) or non-permeabilizing buffer (PBTG without Triton X-100) for 15 min. Samples were incubated with the selected primary antibody diluted in 1% BSA in PBS for 1 h at room temperature, washed with PBS and incubated with the corresponding secondary antibody for 30 min. Wheat germ agglutinin coupled with Alexa fluor 488 was added together with the secondary antibody. DAPI was used for DNA staining. Samples were then mounted in Prolong Gold anti fade reagent (Invitrogen) and cells observed with a Zeiss 710 Laser Scanning Confocal (ZeissCF) microscope.

To swell cells after transfection, cells were incubated in hypotonic media (10% DMEM in H2O, 25 mM HEPES, 50 mM EDTA and 1mM Sodium orthovanadate) for 30 min at 37°C and washed with PBS supplemented with 1 mM Sodium orthovanadate. Cells were then fixed, permeabilized, and incubated with primary and secondary antibodies as described above.

### Cell surface protein biotinylation for SDS-PAGE analysis

Four 100 mm plates of CHO cells were transfected with 15 μg DNA of expression plasmids directing expression of each full-length RTK full and cell-surface biotinylation and isolation of cell surface proteins carried out according to the manufacturer’s instructions (Pierce, Cat # A44390). For biotinylation of the ΔECR variants, an extracellular FLAG tag was added to the N-terminus of each variant.

### Deglycosylation assay

CHO cells were transfected with 1 μg of the indicated expression plasmid, and 8 h post-transfection cells were treated with 2μg/ml Tunicamycin for 20 h. Cells were then treated with 2U/ml PNGase F for 2h at 37 °C, washed, and lysed as previously described.

## Supplemental Figure Legends

**Supplemental Figure 1.**
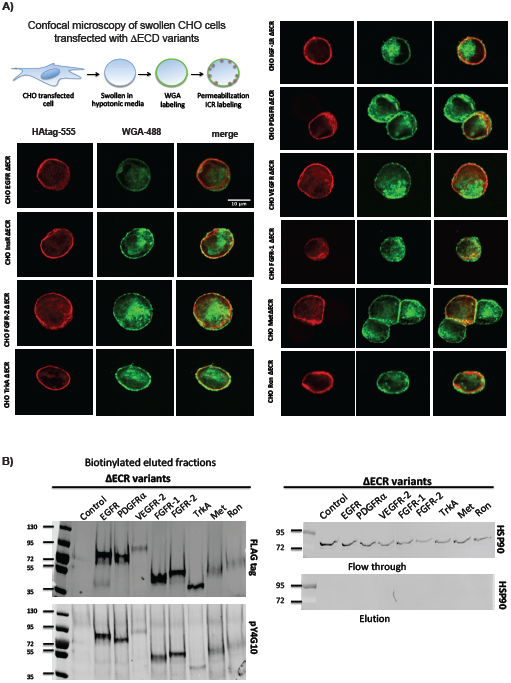
ΔECR forms of all RTKs are trafficked to the cell surface. **A)** Confocal microscopy images of CHO cells transiently transfected with the indicated ΔECR RTK variant showing cell surface expression of each variant. Cells were swollen in hypotonic media, washed, permeabilized and stained with an anti-HA antibody (HAtag-555; red) and wheat germ agglutinin to label the cell surface (WGA-488; green). **B)** N-terminal FLAG-tagged ΔECR variants of the indicated RTKs were transiently expressed in CHO cells. Cell surface proteins were biotinylated, separated by using a streptavidin-affinity matrix, and expressed proteins detected by anti-FLAG Western blot (left). Anti-HSP90 Western blot of streptavidin-affinity Elution and Flow Through fractions showing that cytoplasmic proteins were not detectably biotinylated.

**Supplemental Figure 2.**
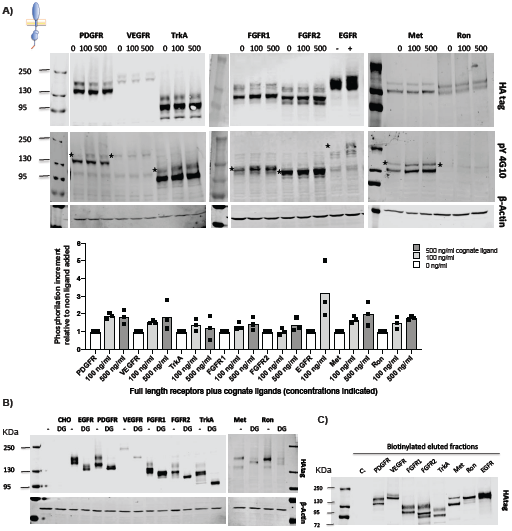
Cell-surface expression, glycosylation and ligand-dependent phosphorylation of native RTKs. **A)** HA-tagged forms of the indicated RTKs were transiently expressed in CHO cells, and expression and phosphorylation levels in the presence and absence of ligand determined by Western blot using anti-HA (HA-tag) and anti-phosphotyrosine (pY4G10) antibodies. Stars indicate slower migrating bands that represent fully processed receptors. Quantification and graphic representation of the band intensities are shown. The phosphorylation intensity was normalized to the expression and the increment was made relative to each receptor without ligand. Dots represent individual values from independent experiments for each sample and the bars represent the calculated mean value of the three experiments. β-Actin was included as a loading control. **B)** Anti-HA Western blot of lysates from CHO cells transiently expressing the indicated RTKs without deglycosylation treatment (-) or treated with Tunicamycin for 20 hours followed by treatment of cell lysates with PNGase F (DG). **C)** Anti-HA Western blot of indicated proteins transiently expressed in CHO cells following biotinylation of cell surface proteins and separation using a streptavidin-affinity matrix.

**Supplemental Figure 3.**
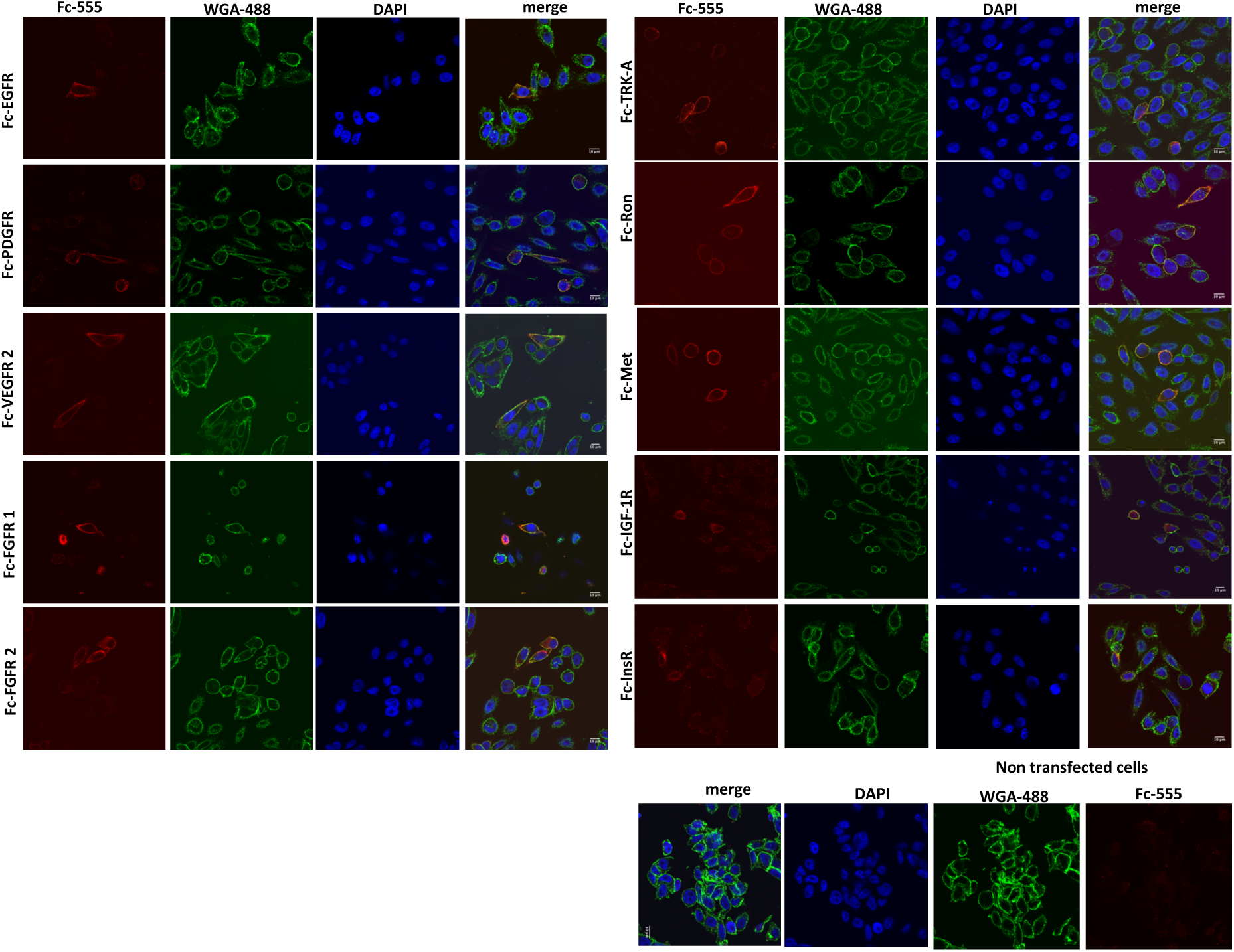
Fc-RTK(TM-ICR) forms of all RTKs are trafficked to the cell surface. Confocal microscopy images of untransfected CHO cells and CHO cells transiently transfected with the indicated Fc-RTK(TM-ICR) variants showing cell surface expression of each variant. Cells were swollen in hypotonic media, washed, permeabilized and stained with an anti-Fc antibody (Anti Fc mouse; red), wheat germ agglutinin to label the cell surface (WGA-488; green), and DAPI (nuclei; blue). The scale bar represents 20 μm.

**Supplemental Figure 4.**
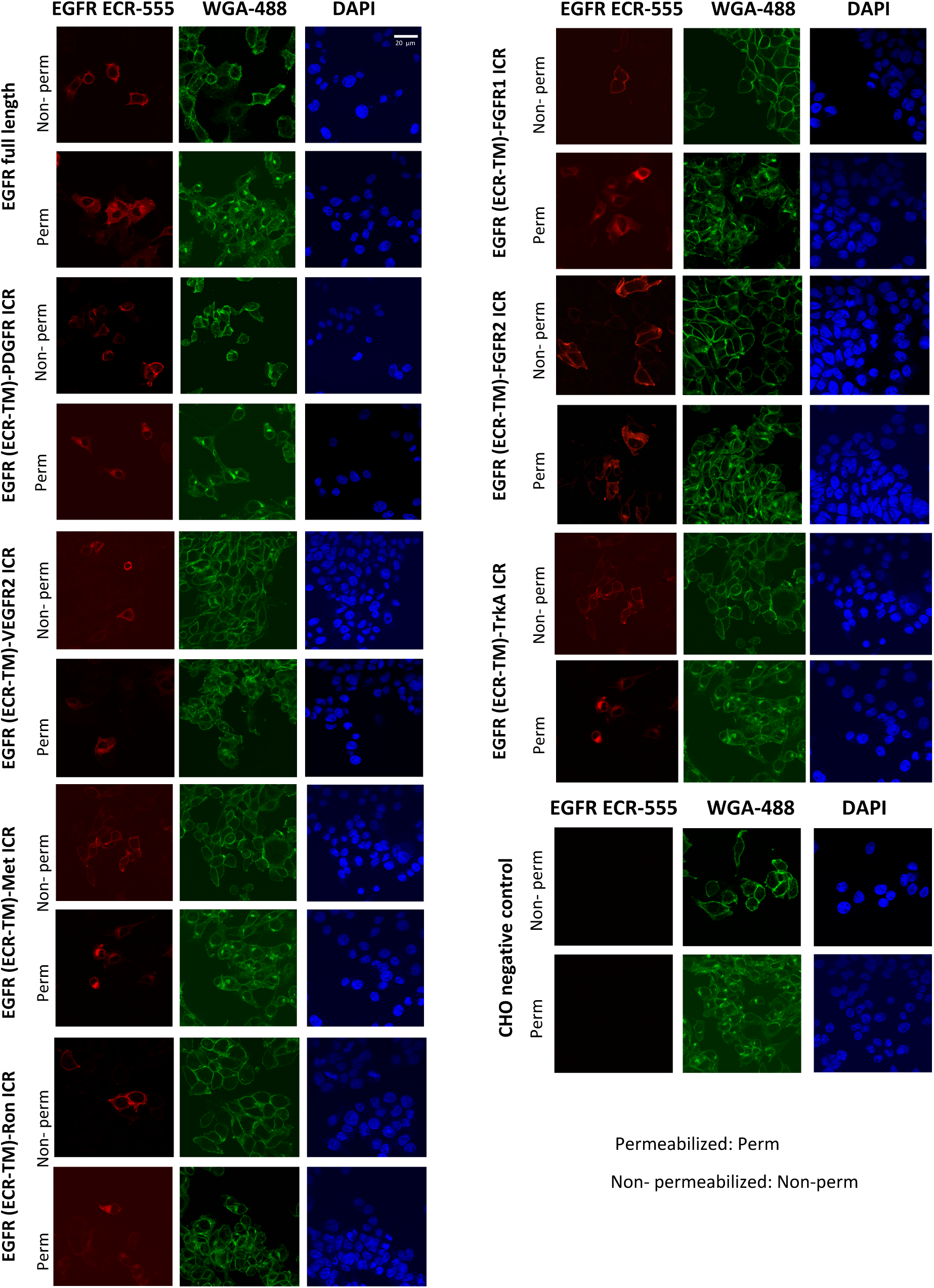
EGFR(ECR-TM)-ICR forms of all RTKs are trafficked to the cell surface. Confocal microscopy images of untransfected CHO cells and CHO cells transiently transfected with the indicated EGFR(ECR-TM)-ICR RTK variant showing cell surface expression of each variant. Cells were swollen in hypotonic media, washed, permeabilized and stained with an anti-EGFR ECR antibody (EGFR ECR; red), wheat germ agglutinin to label the cell surface (WGA; green), and DAPI (nuclei; blue). The scale bar represents 10 μm.

**Supplemental Figure 5.**
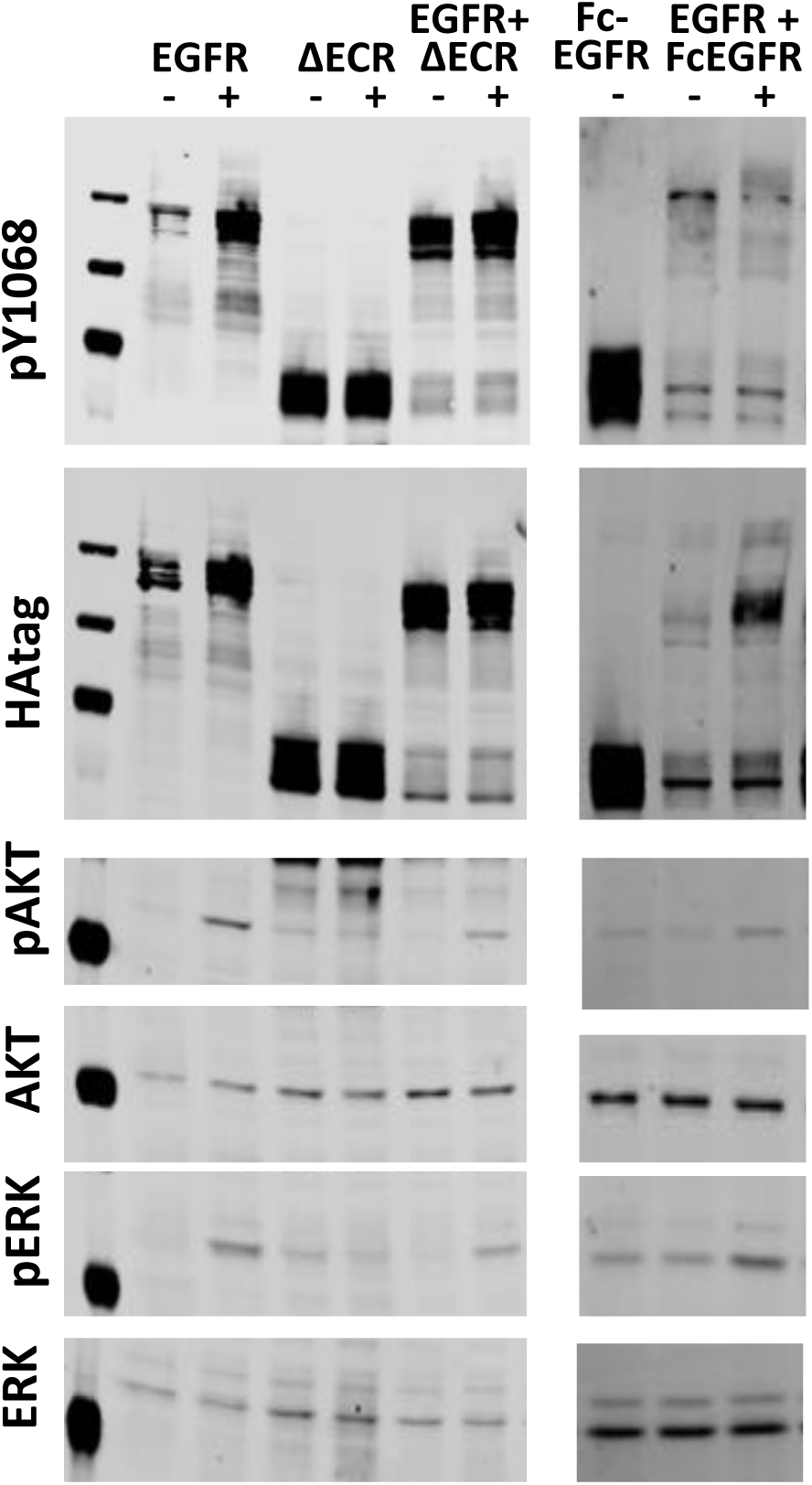
EGFR is able to activate downstream effectors in response to ligand in cells expressing constitutively-phosphorylated Fc-EGFR or ΔECR-EGFR. CHO cells were co-transfected with native EGFR plus Fc-EGFR(TM-ICR) or ΔECR-EGFR, 100 ng/ml EGF were added and lysates were analyzed by Western blot using an anti-HA tag (HAtag), EGFR phospho tyrosine 1068 (pY1068), phospho-Erk 1/2 (pErk1/2), Erk 1/2 expression (Erk 1/2), phospho-Akt (pAkt), and Akt expression (Akt).

## Notes

### Competing Interest Statement

The authors have declared no competing interest.

